# Computationally Grafting an IgE Epitope onto a Scaffold: Implications for a Pan Anti-Allergy Vaccine Design

**DOI:** 10.1101/2020.09.14.296392

**Authors:** Sari Sabban

## Abstract

Allergy is becoming an intensifying disease among the world population, particularly in the developed world. Once allergy develops, sufferers are permanently trapped in a hyper-immune response that makes them sensitive to innocuous substances, such as the protein cupin in peanuts. The immune pathway concerned with developing allergy is the Th_2_ immune pathway where the IgE antibody (targeting innocuous substances) binds to its Fc*ε*RI receptor on Mast and Basophil cells. Currently, there is not a permanent treatment for allergies. This paper discusses a strategy and a protocol that could disrupt the binding between the antibody and its receptor for a potential permanent treatment. Ten proteins were computationally designed to display a human IgE motif very close in proximity to the IgE antibodies’s Fc*ε*RI receptor’s binding site in an effort for these proteins to be used as a vaccine against our own IgE antibody. The motif of interest was the FG motif and it was excised and grafted onto a *Staphylococcus aureus* protein (PDB ID 1YN3). The new structures (motif + scaffold) had their sequences re-designed around the motif to find an amino acid sequence that would fold to the designed structures correctly. These ten computationally designed proteins showed successful folding when simulated using Rosetta’s AbinitioRelax folding simulation and the IgE epitope was clearly displayed in its native three-dimensional structure in all of them. These designed proteins have the potential to be used as a pan anti-allergy vaccine by guiding the immune system towards developing antibodies against the body’s own IgE antibody, thus neutralising it, and presumably permanently shutting down a major aspect of the Th_2_ immune pathway. This work employed *in silico* based methods for designing the proteins and did not include any experimental verifications.

## Introduction

Allergy was first defined by Clemens von Pirquet in 1906 when he discovered that second injections of horse serum caused a severe inflammatory reaction in some, but not all, individuals. He termed this condition Allergy, from the Greek words allos “other” and ergon “works” and therefore the allergy causing agent was called an “allergen” [7]. In the 1960s Kimishige Ishizaka and Teruko Ishizaka demonstrated that allergic reactions are mediated by a new class of antibodies that they discovered and called immunoglobulin E [18] [12], which binds onto a receptor called the high-affinity IgE receptor (Fc*ε*RI) which is found on Mast and Basophil cells and comprises four chains (an *α* extracellular chain with two domains, an intermembrane *β* chain, and two intermembrane *γ* chains protruding into the cytoplasm).

Humans have five antibody types (IgA, IgD, IgE, IgG, and IgM). immunoglobulin G (IgG) is the most abundant antibody type since it targets viral and bacterial pathogens. Immunoglobulin E (IgE) on the other hand, is concerned with parasitic immunity. Since parasites are eukaryotes and closer to other eukaryotes phylogenetically compared to bacteria and viruses, this pathway can target innocuous substances that look like parasites but are not usually harmful, leading to a type of inflammatory reaction termed an allergic reaction, or known medically as type I hypersensitivity.

Thus, IgE antibodies are best known for their role as mediators of the allergic response, which in its most serious manifestations, causes asthma or an anaphylactic shock. Reports of an increase in the number of individuals suffering from allergic manifestations began in the second half of the last century and the incidence of allergy has now reached pandemic proportions [21]. IgE mediated allergic responses have diverse manifestations, which range from mild to severe and can be life threatening. Mammals including humans, dogs, and horses are known to suffer the clinical symptoms of IgE-mediated type I hypersensitivity responses. In spite of extensive worldwide research efforts, no effective active therapeutic intervention strategies are currently available.

One of the perceived reasons for the continual increase in allergy incidence, especially in the developed world, is a hypothesis termed the Hygiene Hypothesis, originally formulated by Strachan [26] [27] [20], it states that a lack of exposure to infectious pathogens in early childhood, i.e. living in an environment too clean, can lead to inadequate immune system development, i.e. a shift from the Th_1_ immune response (bacteria, viruses) to that of the Th_2_ immune response (parasite, allergy), increasing the susceptibility to develop allergy. Further studies in this immunological pathway has shed light into the viability of this hypothesis and showed a correlation between tuberculosis infections in childhood and lack of allergy in adulthood [28].

Currently, the most widely used therapy against allergy is pharmacotherapy, which is a passive immunotherapeutic intervention strategy employing the use of anti-histamines, corticosteroids, or epinephrine, all of which alleviate the symptoms of allergy without curing its underlying cause. The quest to treat allergy is not a new concept, it was first attempted in 1911 [14] when subcutaneous injections of an allergen extract were administered in an effort to desensitise atopic patients to certain allergens. This procedure was successful to treat certain conditions such as anaphylaxis and allergic rhinitis, but it was unsuccessful in treating asthma [17]. This protocol has remained controversial as it has the potential to sensitise patients even more, thus worsening their condition [19]. A novel immunotherapy called sublingual immunotherapy is currently being researched where allergen extracts are given to patients under their tongues [10]. The efficacy of these therapies varies greatly between individuals since doctors do not have a standardised protocol to follow, they usually develop their own protocols according to their own observations and individual successes.

Since allergy incidence have been on the rise globally, a new form of therapy is under development. Though still a passive immunotherapeutic strategy, it employs non-anaphylactogenic antibodies which have demonstrated their capacity to treat type I hypersensitivity responses. These humanised mouse monoclonal antibodies (mAbs), of which Omalizumab [22] is best characterised, are now successful in treating severe forms of allergy, but have been shown to be associated with a number of drawbacks: 1) poor effectiveness in obese patients, 2) logistics and cost, 3) treatment only reduces symptoms temporarily, hence it is a passive immunotherapeutic strategy.

These draw backs logically lead to the potential to develop new active forms of immunotherapeutic strategies, such as a vaccine that primes the immune system against its own IgE antibody, at which point the IgE is neutralised and the allergy disease is terminated. Even though current mainstream research is concentrating on the passive immunisation approach, it is believed that active immunisation is a viable form of treatment against this disease (figure 1).

**Figure 1.**
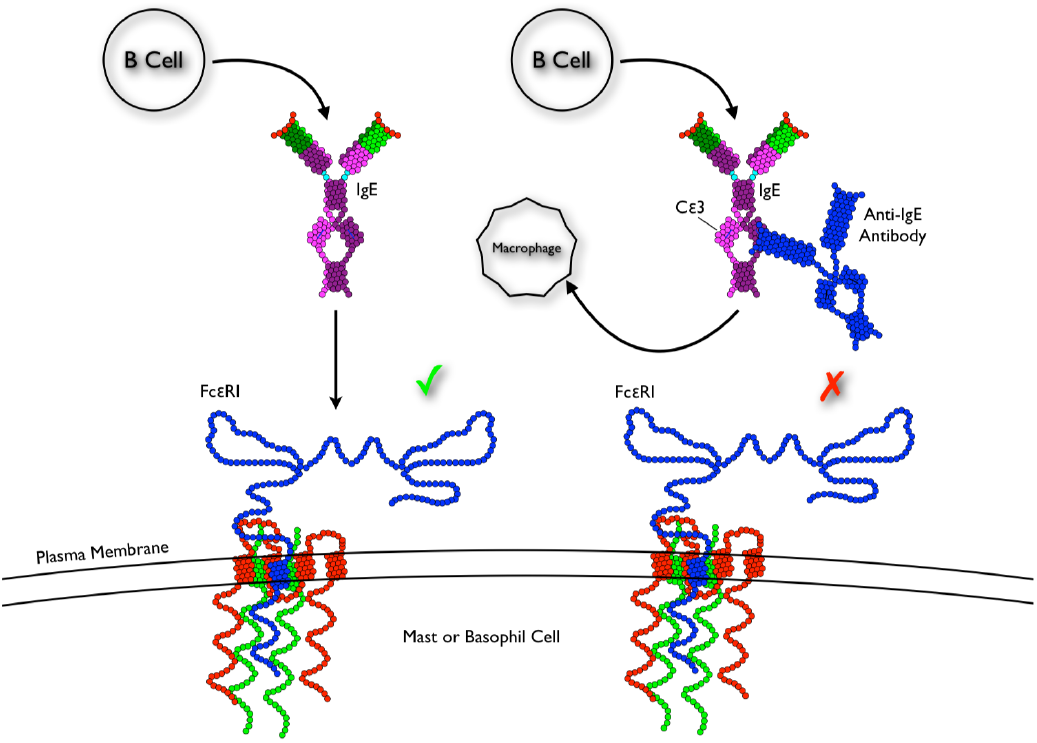
Summery of the pan-anti allergy vaccine therapy concept. Administering a vaccine that is capable of producing antibodies against the body’s own self IgE molecule, and neutralising it by preventing it from binding onto its receptor, could disrupt the entire allergy pathway and potentially curing the disease [30].

This paper proposes a strategy by which a pan-anti allergy vaccine can be computationally designed by excising the mo-tif of interest from the IgE structure and grafting in onto a scaffold protein structure, thus displaying only the motif of interest in its original three dimensional form without any of the surrounding native structure, allowing the immune system to target just that particular motif.

## Methods

The following steps were used to generate a database of scaffold protein structures to search for a an appropriate backbone to graft the motif onto, as well as isolate the IgE motif, graft it onto the found scaffold, then design the scaffold to fold onto the designed protein structure.

### Motif determination and excision

The FG motif (figure 2 purple colour) from the human IgE crystal structure (PDB ID 2Y7Q, chain B, amino acids 420-429 with the sequence VTHPHLPRAL) was chosen due to its very close proximity to the IgE’s receptor binding site (PDB ID 2Y7Q, chain B, amino acids 331-338 with the sequence SNPRGVSA) named the R loop (figure 2 green colour). The FG motif has a ridged structure and it forms a beta sheet loop with an anchoring lysine at position 425 pointing into the core fixing its shape. Since the IgE’s receptor binding site (R loop) was not anchored in place with an amino acid pointing into the core it had a higher degree of movement and thus was not well modelled in the crystal structure, hence it was not chosen (this can be clearly observed when looking at chain C where the same position is missing), furthermore, when the R loop was grafted it assumed multiple structures as can be seen from 2, thus the FG loop was chosen instead. The FG motif was isolated along with the full receptor chain (PDB ID 2Y7Q all of chain A) as separate .pdb files in preparation for grafting. In the 2Y7Q crystal structure only the receptor’s extracellular *α* chain is modeled with its two domains.

**Figure 2.**
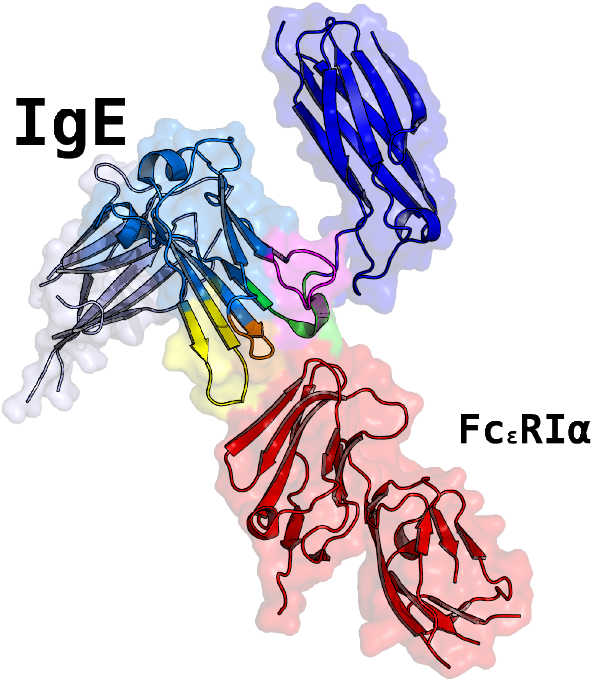
The structure of the human IgE bound to its Fc*ε*RI*α* receptor (2Y7Q). The colours show the different loops that are closet in proximity or forms hydrogen bonds with the receptor when bound. Purple for the FG loop, green for the R loop, orange for the BC loop, and yellow for the DE loop.

### Scaffold database generation

The scaffold database was generated by downloading the entire protein databank using this command:

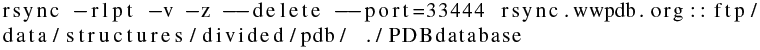

Each structure was unzipped and the original zipped structured deleted to save memory space. Then, each structure with multiple chains was separated into separate .pdb files using this simple python script that uses the biopython [32] python library:

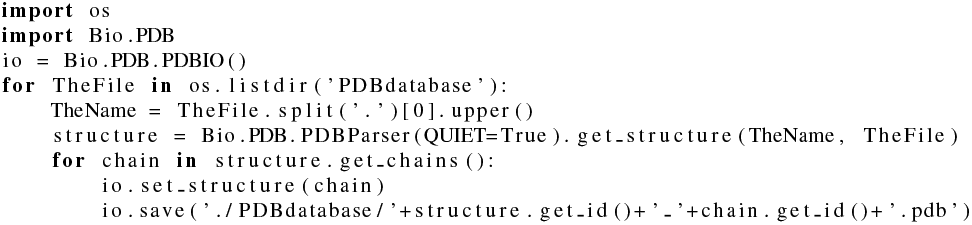

Then desired structures (sizes below 150 amino acids) were isolated using the following bash code:

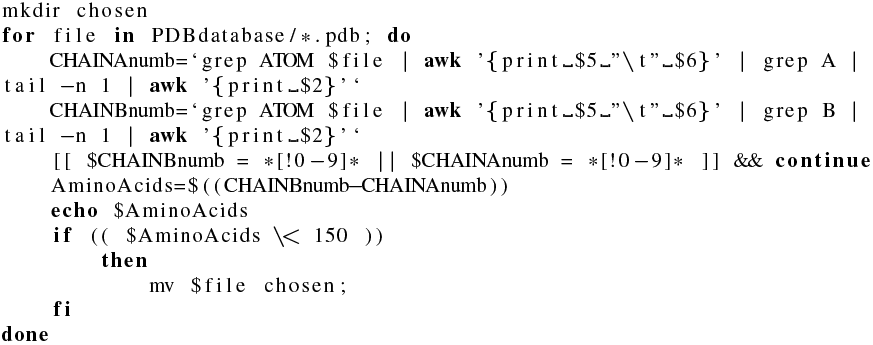

Then, structures were cleaned (removed of any none-peptide atoms) using the following Linux terminal bash command:

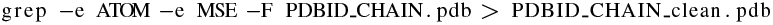

for example:

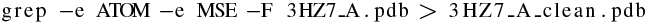

Care must be taken to ensure that the non-canonical MSE (selenomethionine, which is used to solve crystal structures) amino acid is transferred to the cleaned structure which is under the HETATM heading and the MSE amino acid code in .pdb file instead of ATOM heading and the MET amino acid code for methionine. If MSE was not imported it will result in structures with missing selenomethionine. To ensure that the structures are compatible with PyRosetta and will not crash it, they are run through this script (basically just imported then exported):

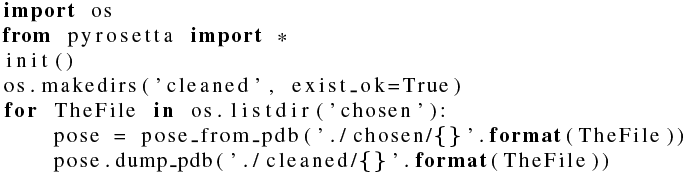

Structures that were not satisfactory were deleted. In this way the scaffold database was constructed.

### Scaffold search and motif grafting

The desired IgE motif (the FG motif) with the sequence VTH-PHLPRAL between positions 420 and 429 in chain B of the protein crystal structure with PDB ID 2Y7Q was isolated along with all of chain A, which was the Fc*ε*RI receptor’s *α* chain since in the 2Y7Q crystal structure only the receptor’s extracellular *α* chain was modeled with its two domains. A scaffold search was performed where the motif was grafted onto each structure within the scaffold database using the epitope grafting protocol [3] [2]. If there was a match within an RMSD value of 1.0 Å or less the grafted structure (with the motif replacing the original backbone on the scaffold) was measured for its clash with the receptor (i.e: to make sure the backbone was not grafted inward or was buried within the structure). If there was no clash with the receptor structure, then the final grafted structure was exported. The code used for this step can be found in this GitHub repository.

### Selective fixed-backbone sequence design

The final grafted structure was tested for folding (in the next section), which failed to converge into a low root-mean-square deviation (RMSD) low free energy score. Thus, to find a sequence that would allow the grafted structure to fold into the desired structure it had to be sequence designed, i.e: find a sequence that would fold into the desired structure. Initially, this was attempted manually by human guided mutations where amino acids were mutated at a strategic location chosen visually to fill in core voids using only amino acids that were specific to the secondary structures of the mutation sites and their layer position calculated by each amino acid’s surface accessible surface area (SASA) using the same criteria in [33]. After several failed attempts, the RosettaDesign fixed-backbone design protocol was employed [13] [8] [1] [11] [15]. The side chains (amino acid identities) of the structure were stochastically mutated and packed using a rotamer library to find the lowest energy structure that would fold into the designed structure. In this protocol, the REF2015 energy function weights were changed to include aa rep 1.0, aspartimid penalty 1.0, buried unsatisfied penalty 1.0, and approximate buried unsat penalt 5.0, which assisted in designing an adequate sequence that fits the backbone structure and increased the energy gap between the desired structure and any other possible undesired fold. The code used for this step can be found in this GitHub repository.

### Folding simulation

To get insight into whether or not the design process was successful, the folding of the sequence-designed-grafted-structure was simulated using the Rosetta [16] AbinitioRelax protocol [23] [6] [4] [5] [24] [25], which employs a Monte Carlo method, where the amino acid sequence is used to construct a straight primary structure, then 3-mer and 9-mer fragments are randomly inserted, the fragments were generated from the Robetta fragment server (http://robetta.bakerlab.org/fragmentsubmit.jsp) using the amino acid sequence. These fragments are backbone torsion angles of secondary structures that were statistically calculated from the amino acid sequence and they help speed up the simulation. Then the structure is randomly moved (backbone and side chain torion angles changed) and its free energy calculated using the REF2015 scoring function which employs first physical principles and some statistical weights [31] using the following equation (details are explained in the original paper):

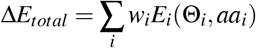

After several cycles of moving and scoring the final structure was exported. This was repeated 1 million times, which results in 1 million simulated structures. These structures are then plotted on a score vs RMSD plot to show how similar they are to the original designed structure. A successful simulation would result in a funnel-shaped plot, where the lowest scoring structures (lowest free calculated energy) results in structures close to the designed structure (low RMSD) since it is assumed that any protein structure resides in the global free energy minima. The code used for this step can be found in this GitHub repository.

## Results

Analysis of several motif positions revealed that the R loop and the FG loop from the human IgE (PDB ID 2Y7Q) are the best candidates for a targeted vaccine due to their proximity to the binding site on the *α* chain of the Fc*ε*RI receptor (figure 2). After several attempts at grafting and designing the R loop onto a *Mycobacterium smegmatis* EsxGH protein (PDB ID 3Q4H replacing the sequence QGDTGMTY at positions 44-51), the FG loop appeared to be the better choice, this was due to the FG loop having an inward pointing leucine, resulting in a ridged loop structure, compared with the R loop that had a high degree of angle freedom, which resulted in a wide range of different structures when grafted, see figure 3 for a comparison between the grafted and designed motif coloured in purple.

**Figure 3.**
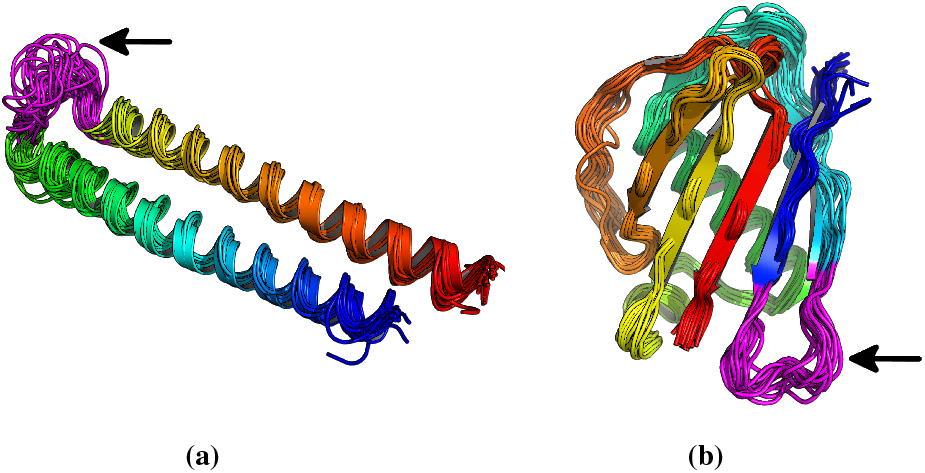
Comparison of grafting the R motif to grafting the FG motif. **A:** Folding simulation of the R loop motif (in purple) after successfully being grafted onto the 3Q4H protein and sequence designed, here showing a large variability of the motif backbone since it lacked an anchor (average RMSD =1.29 Å to the natives motif). **B:** Folding simulation of the FG loop motif (in purple) after successfully being grafted onto the 1YN3 protein and sequence designed, here showing better motif stability (average RMSD = 0.62 Å to the natives motif). Structures rendered through PyMOL [29]

The scaffold search resulted in the FG loop motif being grafted onto a *Staphylococcus aureus* EAP protein (PDB ID 1YN3) [9] structure as well as several other structures (figure 4). The 1YN3 structure was chosen because it had a backbone that was easily simulated by forward folding using the AbinitioRelax protocol (figure 5) when tested as a control on the original wild type crystal structure. Another reason was that the 1YN3 protein is a *Staphylococcus aureus* protein which was expressed in *Escherichia coli* when it was crystallised and is highly antigenic, thus it is predicted to easily crystallise for final structural evaluation and the backbone could instill a strong immune response, which is required to develop antibodies that would bind to the IgE at a stronger affinity than the IgE binds onto its receptor. The motif was grafted between positions 164 and 173 on the 1YN3 structure replacing the sequence ITVNGTSQNI with VTHPHLPRAL (figure 6). As predicted the freshly grafted structure failed a forward fold simulation using AbinitioRelax, this was due to the addition of the motif backbone and side chains severely disrupted the stability of the entire structure. To overcome this, the entire structure was sequence designed by changing and optimising the side chains (except for the motif) while fixing the backbone to stabilise the structure and accommodate the new motif backbone and side chains. At first, manual sequence design was performed, which proved fatal, then the RosettaDesign protocol was successfully used as described in the methods section. Since a failure rate exists between a successful forward fold and a successful crystal structure, the sequence design step was repeated ten times, this resulted in ten structures with the same motif and backbone but different sequences, all of which had a successful forward folding simulation (figure 7). This should increase the probability of synthesising a correctly folded vaccine structure since only one of these structures must pass a crystallography evaluation to be tested as a potential vaccine. If several structures do pass the crystallography evaluation, one structure can be used as a vaccine, while the others are used as a boosts.

**Figure 4.**
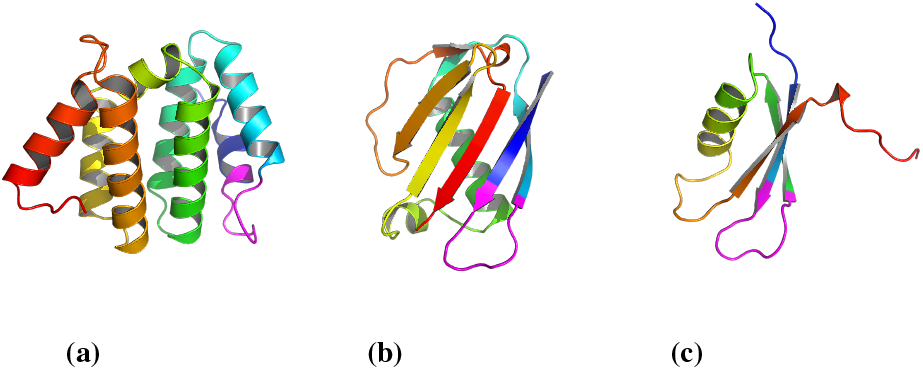
Grafting the FG motif onto three different scaffolds. This figure is showing three structures that successfully accepted the grafted the FG loop **A:** A domain of the human STAM1 VHS (PDB ID 3LDZ chain A) where the ATSEMNTAED sequence at positions 16-25 was replaced by the FG motif. **B:** An EAP domain protein from *Staphylococcus aureus* (PDB ID 1YN3 chain A) where the sequence ITVNGTSQNI at positions 164-173 was replaced by the FG sequence. **C:** The PAAB subunit of the Phenylacetate-CoA Oxygenage from *Ralstonia eutropha* (PDB ID 3EGR chain B) where the VRSKQGLEHK sequence at positions 13-22 was replaced by the FG motif. 1YN3 was chosen since the native structure was easily forward folded using AbinitioRelax (figure 5). Thus it was easier to redesign this structure and it did result is a structure with a large energy gap between the desired structure and any other potential structure.

**Figure 5.**
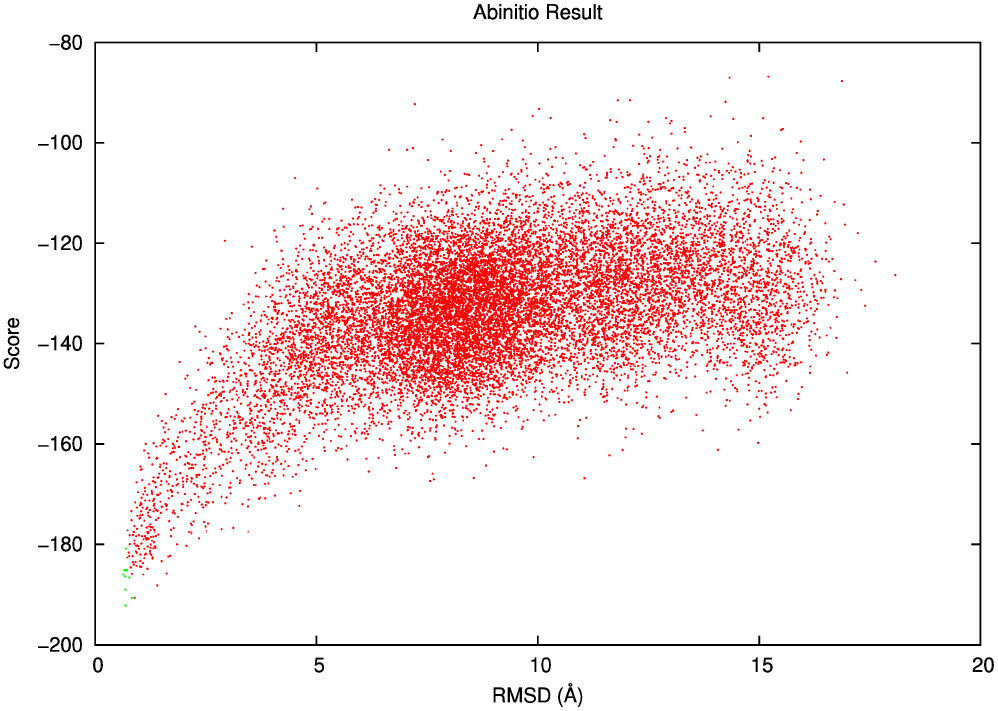
Folding simulation of the native 1YN3 protein structure. AbinitioRelax result of the native 1YN3 protein showing a successful simulation, a funnel shaped plot with the lowest simulated energy close to the predicted energy and RMSD of the structure.

**Figure 6.**
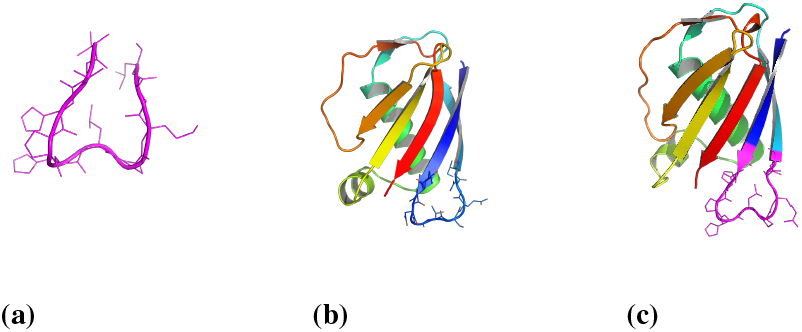
Stages of the grafting protocol. **A:** The native structure of the motif. **B:** The native structure of the *Staphylococcus aureus* EAP protein (PDB ID 1YN3) used here as a scaffold **C:** The final structure after the motif in purple was grafted onto the scaffold, then only the scaffold sequence designed (not the motif).

**Figure 7.**
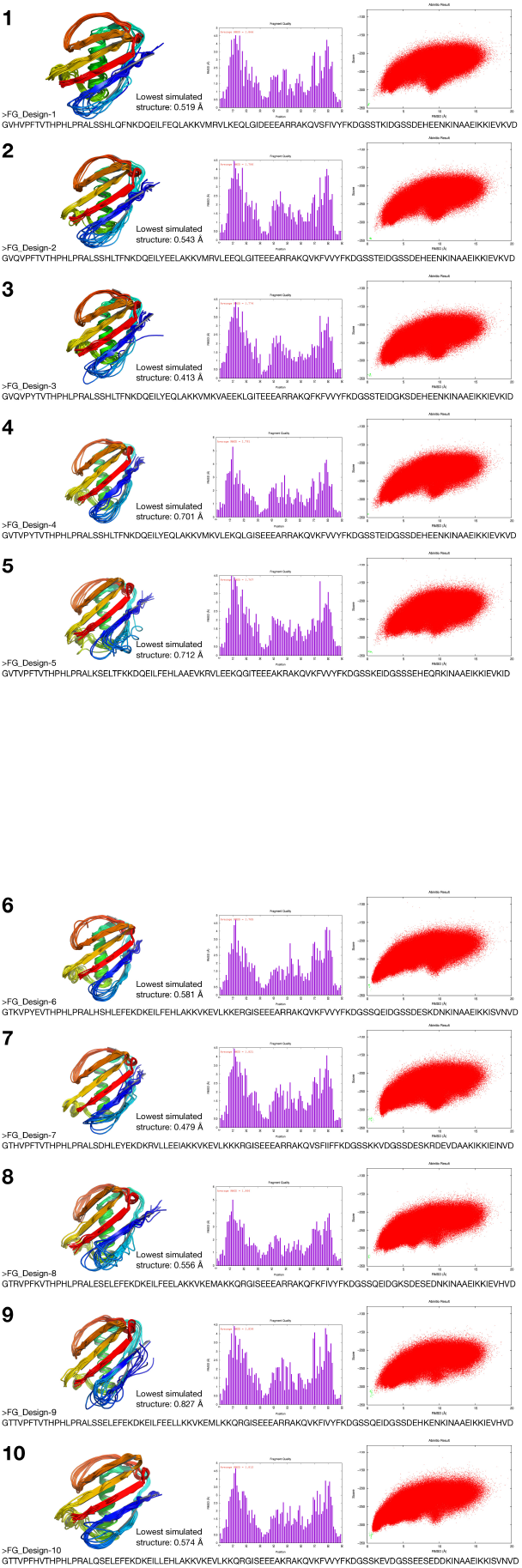
Final designed structures. Ten successfully designed structures that display the FG loop in its native three dimensional structure. The figure shows each designed structure (cartoon) superimposed onto the lowest energy and lowest RMSD structures from the AbinitioRelax simulation (wire) and the corresponding lowest RMSD value of the simulation, thus all structures were predicted to fold within a sub angstrom level of the designed structure giving high confidence that the proteins will have this fold when they are physically synthesised. Also showing are the FASTA sequences of each structure, the fragment quality used in each AbinitioRelax simulation, and the AbinitioRelax plot showing a successful funnel shaped plot for all structures. The green points in each folding simulation are the REF2015 (Rosetta Energy Function 2015) energy score values of the corresponding computationally designed structure after being relaxed thus indicating the lowest possible energy score for each structure and is thus used as a baseline to show were the global minima could be.

The following are the sequences of all the structures, aligned with each other to highlight the differences:

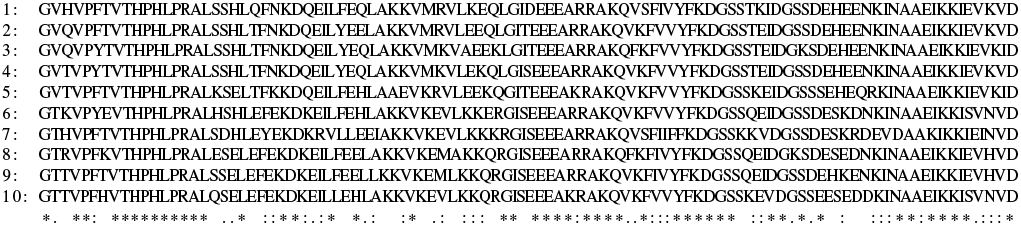

To analyse the structures further, all their FASTA sequences were used to predict the secondary structures of the final proteins. The following are the predicted secondary structures using PSIPRED [34] (H for helix, E for Strand, and C for Coil), *des* is for the designed structure’s secondary structures and *pre* is for predicted secondary structures from the designed structure’s amino acid FASTA sequence:

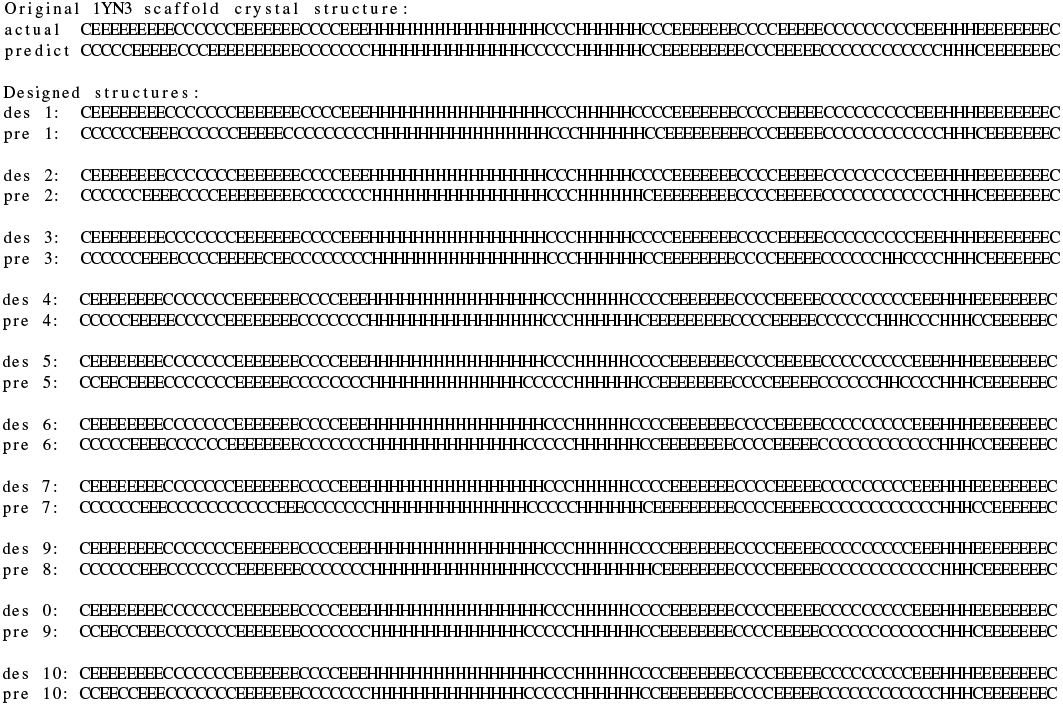

Furthermore, the SWISS-MODEL tool was use to predict the structure of the designed structure from their FASTA sequence as a way to further evaluate their structures computationally. All proteins were predicted to fold as their designed structures 8.

All structures were predicted to fold within a sub angstrom level of the designed structure, giving high confidence that these will be the structures of the proteins when physically synthesised. Each structure must be crystallised and confirmed the correct fold of the protein and the motif before they can be tested on animals.

Molecular Dynamics simulations [36] [37] [38] [39] [40][41][42] [43] were performed on all structures to test the stability of the folded designed structures. Initially the simulation was performed at 300°K using a 0.002 femtosecond time step for 100 nanoseconds (figure 9), all the structures showed an RMSD value around 2 Å. Structures 3, 4, 5, 6, and 8 showed the highest stability (RMSD values mostly less than 2 Å) and a radius of gyration value less than 13 Å. while structures 7, 9, and 10 showed the lowest stability (RMSD values reaching above 2 Å but less than 3 Å at the end of the simulation). The radius of gyration for all structures was around 13 Å. Then the simulation was performed at 400°K using the same parameters to test if the structures would unfold (figure 10). The structures showed less RMSD stability (fluctuating up to 4 Å and 5 Å). Structures 1, 2, 3, 4, 6, 7, 8, 9, and 10 showed low stability by reaching higher RMSD values than the 300°K simulation, while structure 5 showed the highest stability by maintaining an RMSD value between 2 Å and 3 Å. The radius of gyration for all structures remained at around 13 Å but with more variablity than the 300°K simulation.

These simulations can be compaired to the simulation of the original 1YN3 scaffold crystal structure from the Protein Databank (figure 11), where the structure was simulated at 300°K and 400°K using the same parameters. From the simulation at 300°K the structure showed a stable RMSD values (around 2 Å) with a value under 2 Å at the end of the sim-ulation, and a radius of gyration value around 13 Å. While the simulation at 400°K the structure showed a less stable structure with less stable RMSD values (above 2 Å sometimes reaching 4 Å) with a value above 2 Å at the end of the simulation, and a less stable radius of gyration value above 13 Å.

**Figure 8.**
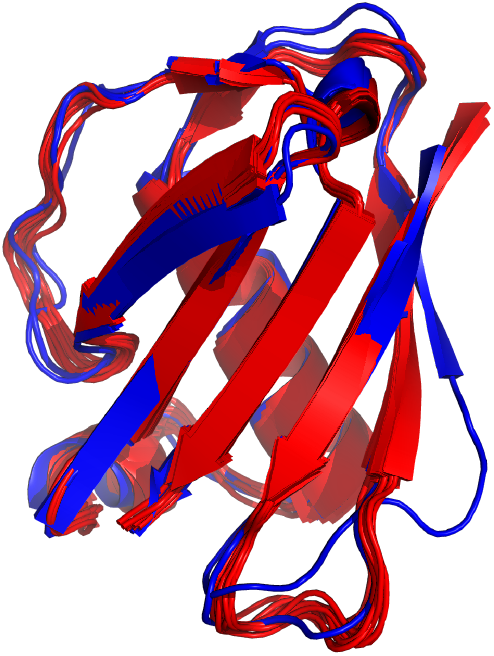
Swiss Model Predictions. The FASTA sequence of each of the designed proteins was used to predict their structure using Swiss Model [34]. Here it can be seen that the predictions from the FASTA sequence predicts similar structures to the desined structures.

**Figure 9.**
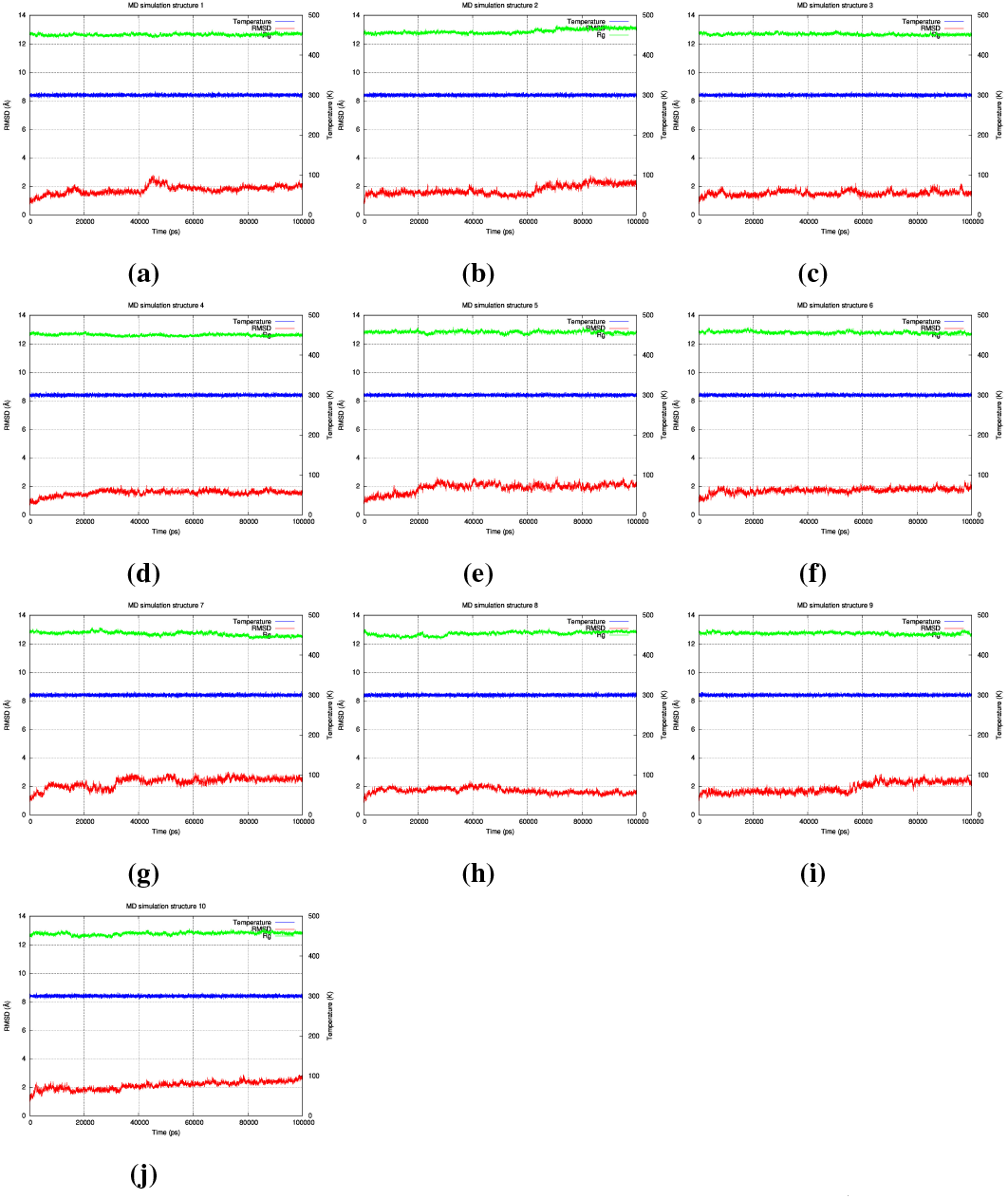
Molecular dynamics simulation at 300°K. Molecular dynamics simulation of all the 10 designed structures at 300°K (26.85°C) for 100 ns (100,000 ps), all the structures showed a stable RMSD value (around 2 Å) where structures 3, 4, 6, and 8 showed the highest stability while structures 7, 9, and 10 showed the lowest stability. The radius of gyration for all structures was also stable (around 13 Å.

**Figure 10.**
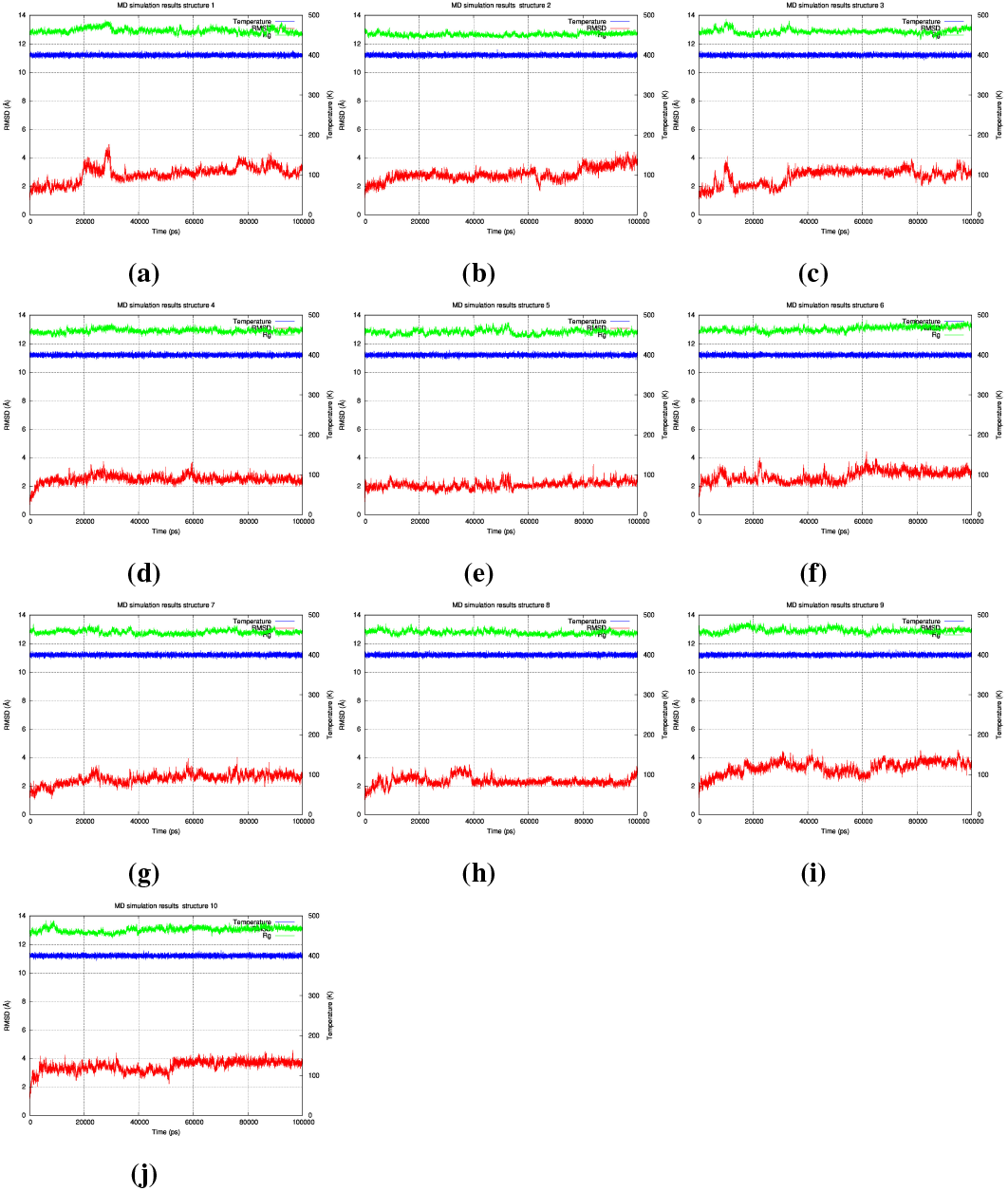
Molecular dynamics simulation at 400°K. Molecular dynamics simulation of all the 10 designed structures at 400°K (126.85°C) for 100 ns (100,000 ps), all the structures showed less RMSD stability (fluctuating up to 4 Å and 5 Å). Structures 1, 2, 3, 4, 6, 7, 8, 9, and 10 showed low stability, reaching high RMSD values, while structures 5 showed the highest stability maintained at 2 Å on average (occasionally reaching 3 Å). The radius of gyration for all structures remained at around 13 Å.

**Figure 11.**
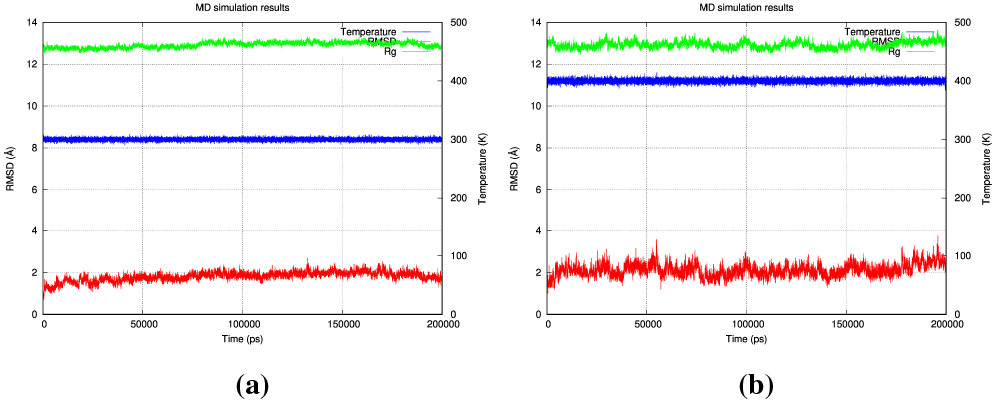
Molecular dynamics simulation of the original 1YN3 crystal structure structure at 300°K and 400°K. Molecular dynamics simulation of the original 1YN3 structure from the Protein Databank at 300°K (26.85°C) and at 400°K (126.85°C) for 100 ns (100,000 ps). **A:** The 300°K simulation showed a structure stable at RMSD value around 2 Å with a final RMSD value under 2 Å at the end of the simulation, it also showed a stable radius of gyration value around 13 Å at the end of the simulation. **B:** The 400°K simulation showed a less stable structure reaching an RMSD value of 3 Å and sometimes coming close to 4 Å with a final RMSD value above 2 Å at the end of the simulation, it also showed a less stable radius of gyration value reaching above 13 Å at the end of the simulation.

It can thus be argued that structures 3, 4, 5, 6, and 8 are the most stable structures at 300°K (26.85°C), while structure 5 is the most stable structures at 400°K (126.85°C).

## Conclusion

This paper describes the protocol for computationally designing proteins that correctly display the three-dimensional structure of the FG strategic motif of the human IgE molecule. The motif was grafted onto the *Staphylococcus aureus* EAP protein (PDB ID 1YN3), which was used as a scaffold structure, then the scaffold/motif was sequence designed multiple times resulting in ten structures each with the same backbone, displaying the same motif in almost its native structure, yet each structure having a different sequence around the motif. Therefore, opening the possibility of using such protein structures as a vaccine and boosts against our own IgE to permanently shut down the allergy pathway regardless of the offending allergen (a pan-anti allergy vaccine). The resulting structures showed agreement in their final folds when simulated with the Rosetta AbinitioRelax folding algorithm, folding to sub angstrom levels when computationally folded from their amino acid sequence’s primary structure. Nevertheless, the only definitive way to determine their realistic physical folds is to solve their structures through X-Ray crystallography or NMR. Furthermore, the efficacy of the proteins in pushing the immune system into developing antibodies against our own IgE at a higher binding affinity than the IgE/Fc*ε*RI receptor’s binding affinity could not be computationally simulated, and thus must be tested on animals to reach a definitive answer. The script that was used to design these proteins is available at this GitHub repository, which includes an extensive README file and a video that explains how to use it.

This work performed initial testing of the hypothesis by employing *in silico* based methods for designing the proteins and did not include any experimental verifications. As a follow up, experimental verifications are required to further test this hypothesis which should include synthesis and purification of all the proteins in a bacterial host (the sequence of each protein is provided in figure 7), testing for binding between the synthesised proteins and known anti-IgE antibodies using the enzyme-linked immunosorbent assay (ELISA), modeling of all the structures through X-ray crystallography to ensure the FG loop is in the correct structure, and finally challanging animals for an immune reaction then testing their sera for binding to the proteins and the human IgE through ELISA, measuring the binding affinity of the antibodies to the proteins and human IgE through Surface Plasmon Resonance (SPR), and testing for IgE/Fc*ε*RI complex distruption through a cell-based mediator release assay [30].

## Availability of data and materials

The code used in this project is available at this GitHub repository which includes an extensive README file and a video that explains how to use the script.

### Grant information

The authors declared that no grants were involved in supporting this work.

### Competing interests

The author has used these results to apply for a patent (filed by the institute) under application numberUS 16/988,076. The patent application is pending.

## Acknowledgements

The corresponding author would like to thank the High Performance Computing Center at King Abdulaziz University for making available the Aziz high performance computer where the corresponding author was able to perform the Epitope Grafting search, the AbinitioRelax folding simulations, and the molecular dynamics simulations.

## Notes

https://sarisabban.github.io/VaxDesign/

### Summary of Updates

* General revision of the manuscript was completed. * Sequence alignment was added to the manuscript. * Molecular Dynamics simulations were performed and added to the manuscript.

